# Nrf2 activator-encapsulating polymeric nanoparticles and LDL-like nanoparticles target atherosclerotic plaque

**DOI:** 10.1101/2020.06.10.144451

**Authors:** Sophie Maiocchi, Sydney Thai, Nicholas Buglak, Ana Cartaya, Arnida Anwar, Ian Corbin, Edward Bahnson

**Affiliations:** Department of Surgery, Division of Vascular Surgery. Center for Nanotechnology in Drug Delivery. MacAllister Heart Institute. University of North Carolina at Chapel Hill. Chapel Hill, NC 27599; Curriculum in Toxicology and Environmental Medicine. University of North Carolina at Chapel Hill. Chapel Hill, NC 27599; Department of Pharmacology. University of North Carolina at Chapel Hill. Chapel Hill, NC 27599; Advanced Imaging Research Center, University of Texas Southwestern Medical Center at Dallas, Dallas, TX 75390, USA; Internal Medicine Division of Liver and Digestive Diseases; RadiologyUniversity of Texas Southwestern Medical Center at Dallas, Dallas, TX 75390, USA; Department of Cell Biology & Physiology. University of North Carolina at Chapel Hill. Chapel Hill, NC 27599

## Abstract

Atherosclerotic vascular disease is the leading cause of death world-wide with few novel therapies available in spite of the ongoing health burden. Oxidative stress is a well-established driver of atherosclerotic progression; however the clinical translation of redox-based therapies is lacking. One of the challenges facing redox-based therapies is their targeted delivery to cellular domains of redox dysregulation. In the current study we sought to develop NPs encapsulating redox-based interventions that exploit passive means of targeting to selectively accumulate in atherosclerotic plaque with the aim of enhancing the intra-plaque bioavailability of interventions. Herein we present two types of nanoparticles (NPs): (i) We have employed flash nanoprecipitation to synthesize polymeric NPs encapsulating the hydrophobic Nrf2 activator drug, CDDO-Methyl, (ii) we have generated LDL-like NPs encapsulating the anti-inflammatory compound, oleic acid (OA). Nrf2-activators are a promising class of redox-active drug molecules whereby activation of Nrf2 results in the expression of several antioxidant and cyto-protective enzymes. Moreover, local activation of Nrf2 within the atherosclerotic plaque can be athero-protective. In this study we characterize the physiochemical properties of these NPs as well as confirm in vitro association of NPs with murine macrophages. In vitro drug release of CDDO-Me from polymeric NPs was determined by Nrf2-ARE-driven GFP fluorescence. In vivo localization was assessed through immunofluorescence of histological sections as well as whole-tissue light sheet fluorescence microscopy. We show that CDDO-Me-NPs and LDL-OA-NPs selectively accumulate in atherosclerotic plaque of two widely-used murine models of atherosclerosis: ApoE^-/-^ and LDLr^-/-^ mice. Overall, these studies underline that targeting of atherosclerotic plaque is an effective means to enhance delivery of redox-based interventions. Future work will assess the therapeutic efficacy of intra-plaque Nrf2 activation or anti-inflammatory actions with CDDO-Me-NPs or LDL-OA-NPs, respectively.

## 1. Introduction

Arteriosclerotic cardiovascular disease (CVD) is the most common cause of death in the entire world according to the World Health Organization. Despite the massive ongoing burden of CVD, since 2000, only 1 out of 10 new US FDA-approved drugs were indicated for CVD and only 1/3 of these had a novel mechanism of action^1, 2^. Current treatments for atherosclerosis involve risk factor management and invasive surgical procedures that carry risks of restenosis and thrombosis^3^. There is a pressing need to develop alternative therapeutic strategies to treat atherosclerotic plaque progression.

There is great interest in the activation of the transcription factor Nrf2 as a mechanistic approach to limit oxidative stress and inflammation associated with atherosclerosis^4, 5^. Nrf2 is a master regulator of the cellular response to oxidative or electrophilic stress through the transcription of numerous cytoprotective and antioxidant genes including heme oxygenase 1, superoxide dismutase 1 and catalase. Under homeostatic conditions, Nrf2 is subject to proteasomal degradation, however upon oxidative or electrophilic stress, inhibition of the ubiquitination process takes place, resulting in Nrf2 nuclear translocation where it binds to the Antioxidant Response Element (ARE) and initiates expression of its target cytoprotective genes. Nrf2 and its target genes have been found to have local anti-atherogenic effects in the vascular wall, including endothelial cells, vascular smooth muscle cells, and macrophages^6-9^. Nrf2 also limits inflammation by directly impeding transcription of pro-inflammatory cytokines such as IL-1β and Il-6^8^. In support that therapeutic activation of Nrf2 would be beneficial in the context of atherosclerosis; compounds which are also known as Nrf2-inducers (tBHQ^10^, Ebselen^11^, CDDO-Me analogue dh404^12^, and oleanolic acid^13^) augmented endogenous antioxidant systems and limited inflammation to prevent atherosclerosis development or progression in diabetes-aggravated atherosclerosis.

Despite the fact that a considerable body of pre-clinical studies indicate that atherosclerosis is driven by oxidative processes^14, 15^, the clinical translational of redox-based therapies has been lacking. One of the greatest challenges related to therapeutically targeting redox dysregulation in CVD, is the specific delivery of redox-based interventions to cellular domains where redox dysregulation is occurring^15^. Targeted nanomedicine is a promising approach to counter this challenge in CVD^5, 16-18^.

Herein we report the encapsulation of the potent Nrf2 activator drug, CDDO-Methyl into polymeric nanoparticles via flash nanoprecipitation^19^. We characterize these nanoparticles for their physiochemical characteristics as well as the *in vitro* release of CDDO-Methyl. Additionally, we report the encapsulation of the anti-inflammatory compound oleic acid in LDL-like nanoparticles^20, 21^. Oleic acid was chosen as a representative loading molecule for bioactive fatty acids and ω-3 PUFAs such as docosahexaenoic acid (DHA), that have been shown to have benefits in cardiovascular disease^22^. We go on to show that both these nanoparticles accumulate in atherosclerotic plaque of athero-prone mice (ApoE^-/-^ and LDLr^-/-^). These studies demonstrate the intra-plaque delivery of antioxidant-based therapeutics.

## 2. Materials and Methods

### 2.1 General materials

CDDO-methyl (Sigma Aldrich, St. Louis, MO. SMB00376-100MG), Synperonic-PE-P84 pluronic tri-block co-polymer (Sigma Aldrich, St. Louis, MO. 713538-1Kg), DMSO (Fisher Scientific, D128-1), platelet derived growth factor–BB (PDGF-BB, Sigma Aldrich, St. Louis, MO. P4056-50UG), DMEM (11885-084; Gibco, Grand Island, NY). DMSO (BP231; Thermo-Fisher Scientific, Waltham, MA), F-12 nutrient mix Ham’s media (11765-054; Gibco), Glucose (50-99-7, Sigma-Aldrich), DMEM High glucose (4.5g/L) (11995-065, Gibco), DMEM low glucose (1g/L) (11885092, Gibco), RPMI 1640 media (11875135, Gibco), Heat-inactivated fetal bovine serum (FBS) (16140071; Gibco), Penicillin-Streptomycin 10,000U/mL (15140122, Gibco), Glutamine (200mM, 25030081; Gibco), Paraformaldehyde (158127; Sigma-Aldrich). PBS (20–134; Apex Bioresearch Products, San Diego, CA). Trypsin-EDTA (0.05%) (25300054; Gibco).

### 2.2 Cell Culture

All cells were cultured in an incubator at 37□°C with 5% CO_2_. RAW 264.7 macrophage cells (ATCC, TIB-71) were cultured in DMEM, high glucose (4.5g/L) supplemented with 10% FBS, 1% Penicillin-streptomycin. Primary murine aortic smooth muscle cells (VSMC) were isolated from the thoracic aorta of 10–12-week male mice. VSMC were used between passages 4–9 for all experiments. VSMC were maintained in low glucose (1 g/L) 1:1 DMEM:F-12 media supplemented with 10% FBS and 1% Penicillin-streptomycin with a final concentration of 4mM glutamine. For assays the cells were synchronized through serum-starvation (24 hours; same media but without the FBS supplement), and treated for assays with media supplemented with 25□ng/mL PDGF-BB.

H1299 cells containing a GFP fragment retrovirally inserted into the second intron of the NQO1 gene (130207PL1G9) were a gift from Tigist Yibeltal and the Major Lab (UNC, LCCC), Uri Alon and the Kahn Protein Dynamics group^23, 24^. These cells were cultured in RPMI 1640 media supplemented with 10% FBS and 1% Penicillin-streptomycin. CDDO-Me was stored in stock concentrations in DMSO at −□20□°C and diluted directly in media for treatment. Equal volume DMSO was included for all controls. CDDO-Me-NPs were stored in PBS and diluted into media. Equal volumes of PBS and equal mass concentrations of polymer were included as a control.

### 2.3 Synthesis of polymeric CDDO-Me containing nanoparticles (CDDO-Me-NPs)

Polymeric CDDO-Me NPs were synthesized via flash nanoprecipitation (FNP) with a confined impinging jet mixer (CIJ) as previously described^19^. Briefly, CDDO-Me and Synperonic PE-P84 were dissolved in tetrahydrofuran (THF, Fisher Scientific, T425-1) at 5 mg/mL each. PBS was used as the aqueous solvent. The two solvent streams were mixed together with the CIJ into 4 mL of PBS. This suspension was then dialyzed overnight against PBS. To generate fluorescent nanoparticles, DiD (Fisher Scientific, D7757, final concentration 0.25mg/mL) was added to the organic solvent prior to injecting the solvent streams through the CIJ.

### 2.4 Preparation of LDL-oleic acid-nanoparticles (LDL-OA-NPs)

Human LDL was isolated from apheresis plasma of patients with familial hypercholesterolemia using sequential density gradient ultracentrifugation^25^. Incorporation of the lipophilic carbocyanine dye DiD, 1,1′-dioctadecyl-3,3,3′,3′-tetramethylindodicarbocyanine (Thermo Fisher Scientific) and unesterified oleic acid (Nu-chek Prep, Inc) into LDL was performed by the reconstitution method as described previously^26^.

### 2.5 TEM

Conventional TEM images were captured on a JEOL JEM 1230 TEM at 80 kV, LaB6 filament, with a Gatan Orius SC1000 CCD camera and Gatan Microscopy Suite 3.0 software. CDDO-Me-NPs at 0.3 mg/mL in phosphate buffered saline (PBS) were prepared for TEM by pipetting 8 μL atop Formvar on copper 400 mesh TEM grids (Ted Pella) that were treated with glow discharge (PELCO easiGlow^™^ Glow Discharge Cleaning System). After two minutes, samples were rinsed with deionized water and stained with 2% uranyl acetate negative stain for two minutes prior to imaging.

### 2.6 Nanosight nanotracking

The nanoparticle hydrodynamic diameter size via particles/mL distribution was measured using the NanoSight NS500 (Malvern Panalytical, Ltd, UK). Samples were prepared in 10mM PBS at 0.01-0.005mg/mL. At least 5 measurements of 40 s were taken per sample at 25°C.

### 2.7 DLS

The nanoparticle hydrodynamic diameter size via intensity distribution was characterized using a Zetasizer Nano ZS (Malvern Panalytical, Ltd, UK). Samples were prepared in 10mM PBS at 0.05mg/mL. At least 10 scans were taken per measurement, three measurements per sample at 25°C.

### 2.8 HPLC-UV-VIS

CDDO-Me loading capacity and final concentration in nanoparticle suspensions were obtained using an Agilent High Performance Liquid Chromatography (HPLC) system with a UV-VIS detector (Agilent Technology 1200 series). CDDO-Me was resolved with a Supelco Analytical Nucleosil C18 HPLC column (Cat# Z226181, 25 cm x 4.6 mm, 5μm particle size). The area under the peak was recorded for the 200 nm wavelength filter. The mobile phase consisted of A: 0. 1% (v/v) trifluoroacetic acid in Acetonitrile and B: 0. 1% (v/v) trifluoroacetic acid in water, at an isocratic flow of 1 mL/min consisting of 85% A. All mobile phase solvents were HPLC-grade. Samples, 10 μl, were prepared as 5-50μg/mL solutions in 90% HPLC-grade acetonitrile. A standard curve with CDDO-Me was conducted each time.

### 2.9 Nrf2 activation assay

H1299 cells were seeded at 20,000 cells per well in a 96 well black glass-bottom plate (Greiner 96 well plates, 655891, VWR). Media was replaced with media containing treatments (1% DMSO, CDDO-Me (10-200nM), CDDO-Me-NPs (10-200nM) and equivalent concentrations of polymer). 24 hours later, wells were imaged using Gen 5 software (BioTek Instruments, Winooski, VT) on a Cytation 5 plate reader (BioTek Instruments) (37°C, 5% CO_2_) with a GFP filter cube (BioTek Instruments, part #:1225101) and a Texas Red filter cube (BioTek Instruments: Part # 1225102). Exposure times were fixed for each well. Cells were counted automatically by the Gen 5 software by thresholding in the Texas Red channel.

### 2.10 Macrophage Internalization Assay

RAW 264.7 macrophages were seeded at 500,000 cells per well in a 12-well corning sterile cell culture plate. Cells were incubated overnight in high glucose DMEM (Gibco) supplemented with 10% FBS, 1% penicillin/streptomycin). Cells were washed with HBSS and then treated with fluorescent DiD-loaded CDDO-Me-NPs to a final concentration of 5μg/mL DiD in media or equivalent amounts of PBS. Cells were incubated at either 4°C or 37°C for 18 hr. At the end of the incubation period, cells were washed with cold HBSS and then lysed with cold 1% Triton-X-100 in PBS (1mL per well). Cells were scraped, centrifuged (14,000 rpm, 15 min, 4°C). Cell lysate was measured for its fluorescence (excitation λ = 640nm, emission λ = 690nm) using quartz cuvettes (Fisherbrand by Hellma Quartz Semimicro cells, Cat. No. 14-385-916A) and a spectrofluorometer (Spectra Max M3) and recorded in SoftMax Pro Software. Fluorescence was converted to μg/mL DiD with a standard curve. Cell lysate was measured for protein in μg/mL using the BCA assay (thermo fisher). Results were normalized and expressed as μg of DiD / μg of protein.

### 2.11 Macrophage Association Assay

RAW 264.7 macrophages were seeded at 50,000 cells per well in a 96 well black glass-bottom plate (Greiner 96 well plates, 655891, VWR). Cells had been previously incubated with folic acid deficient RPMI medium (10% FBS, 1% penicillin/streptomycin) for at least 48 hrs prior to seeding for experiments. After seeding they were maintained in folic acid deficient RPMI medium (10% FBS, 1% penicillin/streptomycin). Cells were treated with 10μg/mL Hoescht dye for 30 minutes at 37°C, 5% CO_2_. Cells were then imaged every 4 hours continuously over 18 hours after having media replaced with DMEM no phenol red media (10% FBS, 1% penicillin/streptomycin) containing treatments (PBS (20%) or CDDO-Me-NPs; final concentration of 0.1mg/mL). Alternatively, cells had their media replaced with folic acid deficient RPMI medium (10% FBS, 1% penicillin/streptomycin) containing treatments (PBS (20%) or CDDO-Me-NPs; final concentration of 0.1mg/mL) and were incubated for 18 hours and were imaged just at the 18 hr timepoint. Wells were imaged using Gen 5 software (BioTek Instruments, Winooski, VT) on a Cytation 5 plate reader (BioTek Instruments) (37°C, 5% CO_2_) with a DAPI filter cube (BioTek Instruments, part #:1225100) and a Texas Red filter cube (BioTek Instruments: Part # 1225102). Exposure times were fixed for each well. Cells were counted automatically by the Gen 5 software by thresholding in the DAPI channel.

### 2.12 VSMC migration

VSMC were seeded at 100,000 per well in a 12-well plate. Scratch wound assay was performed on synchronized VSMC in a 12-well plate at 70% confluency and cells were treated with 100-200nM CDDO-Me and CDDO-Me-NPs. Wells were imaged at time of scratch and 24□h and 48 h later using Gen 5 software (BioTek Instruments, Winooski, VT) on a Cytation 5 plate reader (BioTek Instruments). The number of cells that had migrated within the scratch region were quantified using ImageJ software.

### 2.13 Animals and diet

All animal handling and experimental procedures were approved by the Institutional Animal Care and Use Committee at the University of North Carolina – Chapel Hill. (Ref no. ##). Four to six-week-old male apoE^-/-^ mice (B6.129P2-Apoe^tm1Unc^, stock number: 002052), LDLr^-/-^ (B6.129S7-*Ldlr*^*tm1Her*^/J, stock number: 002207) and C57Bl/6 mice (stock number: 000664) were purchased from Jackson laboratory. C57Bl/6 mice were fed standard chow. Mice were allowed ad libitum access to food and water throughout the study. After 1 week acclimation in the Division of Comparative Medicine (DCM) facility, apoE^-/-^ and LDLr^-/-^ were placed on a western high-fat diet containing 40% fat, 17% protein, 43% carbohydrate by kcal and 0.15% cholesterol by weight (RD Western Diet, catalog number: D12079Bi). apoE^-/-^ and LDLr^-/-^ mice were fed with the high-fat diet over the course of 15-19 weeks. Mice underwent experimental procedures and were sacrificed when they were approximately either 19-21 weeks old or 23-25 weeks old.

### 2.14 Tissue processing

Organs (heart, aorta (brachiocephalic tree, arch, thoracic and abdominal), spleen, liver, kidney, lung, muscle tissue (quadriceps), intestinal tissue) were harvested after in situ perfusion-fixation with 10 mL of PBS and 10 mL of cold 2% paraformaldehyde in PBS. Organs were placed in 2% paraformaldehyde in PBS for 2□h at 4□°C. The heart was processed for sectioning of the aortic root as previously described^27^. Briefly, the heart was cut across in a diagonal slice from the left to right atrium and the atria placed in 30% sucrose overnight at 4□°C. Atria were quick-frozen in O.C.T. (4583; Tissue-Tek, Torrance, CA) and stored at −□80□°C. 10□µm sections were cut throughout the entire aortic root and stored at −80°C. The other organs were placed in PBS with 0.05% sodium azide and stored at 4°C for processing for Light Sheet fluorescence Microscopy (LSFM).

### 2.15 Immunofluorescence

Tissue sections were imaged via fluorescence microscopy using an X-Cite® 120 LED Boost (Excelitas Technologies) coupled with a Zeiss Axio Imager A.2 with either a Zeiss Objective Plan-Apochromat 5x/0.16 M27 (FWD=12.1mm), or a Zeiss Objective Plan-Apochromat 20x/0.8 (WD=0.55mm). Fluorescence microscopy images were captured with a Zeiss AxioCam HRm camera and a Zeiss 60N-C 1” 1.0X camera adapter, and observed digitally with AxioVision SE64 Rel. 4.9.1 software. Exposure times and laser strength were fixed when imaging all samples.

### 2.16 Oil Red O staining

Tissue sections were fixed for 10 min with 10% neutral buffered formalin. They were then washed with water for 10 minutes, rinsed with 70% ethyl alcohol for 5 mins. They were then stained for 15 min with freshly prepared Oil Red O working solution (Oil red O solution, Sigma Aldrich, O1391-500mL, dilute 3:2 in deionized H_2_O). Stained slides were then rinsed with 70% ethyl alcohol (5 min), water (30 sec), and stained with hematoxylin (3 min), washed with H_2_O (2 min), then 0.5% acid alcohol solution (5 seconds; 5mL conc. HCl, 1000mL 70% ethanol), then H_2_O (1 min), then 0.5% ammonia water solution (10 sec; 5mL conc. Ammonium hydroxide, 1000 mL water), then finally washed with water. Slides were allowed to dry and then coverslipped and mounted with Aquamount.

### 2.17 Light Sheet Fluorescence microscopy (LSFM)

#### Delipidation and permeabilization

Adipo-clear was performed as previously reported.^28, 29^ Briefly, fixed samples were washed in 20%, 40%, 60%, 80% and 100% methanol/0.1% Triton X-100 (Sigma Aldrich X100-500ml)/0.3 M glycine/0.01% Sodium azide (Sigma Aldrich G2879) buffer (B1N buffer, pH 7.0) for 30 minutes each with shaking at 4°C. Samples were then delipidated and permeabilized with 100% dichloromethane (DCM; Sigma-Aldrich 270997) for 30 minutes, then 1 hr, then 30 minutes. Samples were washed twice with 100% methanol for 30 minutes each, followed by rehydration with a series of 30 minute washes with 80%, 60%, 40% and 20% methanol in H_2_O/B1N buffer. After a final wash in B1N buffer overnight, samples were transferred to PBS/0.1% TritonX-100/0.05% Tween 20/2 μg/mL heparin/0.01% sodium azide buffer (PTxWH buffer, pH 7.4) for 2 hr prior to staining.

#### Immuno-staining

The following primary antibodies were used at the following concentrations: Rabbit polyclonal anti-CD31 (Abcam, 28364, 1:50 dilution); Rat Anti-mouse CD68 (Biorad, MCA1957T, 1:200 dilution, 5μg/mL); Rabbit polyclonal anti-Histone-H2B (Abcam, 1790, 1:200 dilution). The following secondary antibodies were used at the following concentrations: Donkey anti-rabbit AF568 (Invitrogen, A-10042, 10μg/mL, 1:200 dilution); Donkey anti-rabbit AF790 (Invitrogen, A-11374, 10μg/mL, 1:200 dilution); Goat anti-rat PE (Biorad, STAR73, 1:10 dilution). All antibodies were diluted into PTxWH buffer. The antibody dilution was centrifuged briefly at ∼20,000g for 10 min to prevent introducing antibody precipitates or aggregations. The samples were incubated with primary antibody for 4-5 days at room temperature and then washed with PtxWH buffer in a series of incubation steps (5 min, 10 min, 15 min, 20 min, 1 hr, 2 hr, 4 hr, overnight). The samples were then incubated with secondary antibodies for 5-6 days at room temperature. They were then washed again with a similar incubation protocol.

#### Agarose embedding and tissue clearing

Samples were embedded in 1.0% agarose in TAE buffer. Samples were dehydrated in with a series of washed with 25%, 50%, 75% and 100% Methanol/H_2_O for 30 minutes at a time. They were then incubated in 100% DCM for 3 hrs. They were then incubated with 100% dibenzyl ether (DBE) overnight with mild shaking. They were washed with fresh 100% DBE for 2 hr with shaking. They were henceforth stored in 100% DBE at room temperature in the dark prior to imaging.

#### Imaging

Imaging was performed in the Microscopy Services laboratory of UNC. The Microscopy Services Laboratory, Department of Pathology and Laboratory Medicine, is supported in part by P30 CA016086 Cancer Center Core Support Grant to the UNC Lineberger Comprehensive Cancer Center. Imaging was carried out using a LaVision BioTec Ultramicroscope II equipped with zoom body optics, an Andor Neo sCMOS camera, an Olympus MVPLAPO 2X/0.5 objective, and a 5.7-mm working distance corrected dipping cap (total magnifications 4X zoom). Cleared agarose-artery blocks were mounted in a sample holder and submerged in a 100% DBE reservoir. Aorta were imaged at 2X mag (2X zoom), using the three light sheet configuration from a single side, with the horizontal focus centred in the middle of the field of view, an NA of 0.063 (beam waist at horizontal focus = 17□µm), and a light sheet width of 40% (adjusted depending on artery length to ensure even illumination in the *y*-axis). Pixel size was 1.52□µm and spacing of Z slices was 5□µm. Up to four channels were imaged per sample: autofluorescence with 488□nm laser excitation and a Chroma ET525/50m emission filter was used for both species; 561 nm laser excitation and a Chroma ET600/50m emission filter; 647 nm laser excitation and a Chroma ET690/50m emission filter; 785 nm laser excitation and a ET800LP emission filter. The chromatic correction module on the instrument was used to ensure both channels were in focus. To reduce bleaching, while setting up the parameters for image acquisition the sample was viewed with low laser power and long exposures; this was inverted when acquiring data to maximize acquisition speed.

#### Image capturing

LSFM-saved TIFF files (Imspector Pro v5.1.350) were converted to an Imaris (Bitplane, Oxford Instruments) file using Imaris File Converter software (v9.3.1). Acquired LSFM images were first analysed using the 2D slice function and snapshots were obtained of cross-sections in the *x*-*y, x*-*z*, and *y*-*z* axis. The file was cropped to only include the artery to reduce file size and expedite all further steps.

### 2.18 Statistical analysis

Numerical data are represented as means□±□STDEV. Statistical analyses were performed using an unpaired Student’s *t-*test, one-way ANOVA or two-way ANOVA, followed by Tukey’s post-hoc test, as appropriate with a *p*-value□<□0.05 considered statistically significant (OriginLab, Northampton, MA).

## 3. Results

### 3.1 Nanoparticle characterization

Herein we utilized flash nanoprecipitation to generate polymeric nanoparticles encapsulating CDDO-Methyl, a potent nanomolar activator of the Nrf2 transcription factor, as well the lipophilic dye, DiD^19^. We measured their hydrodynamic diameter by dynamic light scattering (DLS) to show that they had an average size of (185.38 ± 42.39) nm (Fig. 1A). DLS also revealed a polydispersity index of 0.271 ± 0.124, demonstrating that they were relatively uniform. We also performed nanosight nanotracking analysis and found a similar average diameter of (186.76 ±83.75) nm (Fig. 1B). We additionally confirmed a spherical shape and uniform size via transmission electron microscopy (TEM) (Fig. 1C,D). Finally, we performed HPLC-UV-VIS analysis of the nanoparticles to measure loading efficiency and loading capacity. Loading efficiency was determined by measuring the amount of CDDO-Me encapsulated in freshly synthesized nanoparticles vs. those that had been dialyzed overnight. Loading capacity was determined by measuring the mol of CDDO-Me present in a solution of solubilized nanoparticles and dividing this by the total mass of CDDO-Me and polymer present. In this manner, the loading efficiency was found to be (87±3)% and a loading capacity of (35±7)%. We additionally found that the nanoparticle size remained unchanged over a 40 hour time period (data not shown).

**Figure 1.**
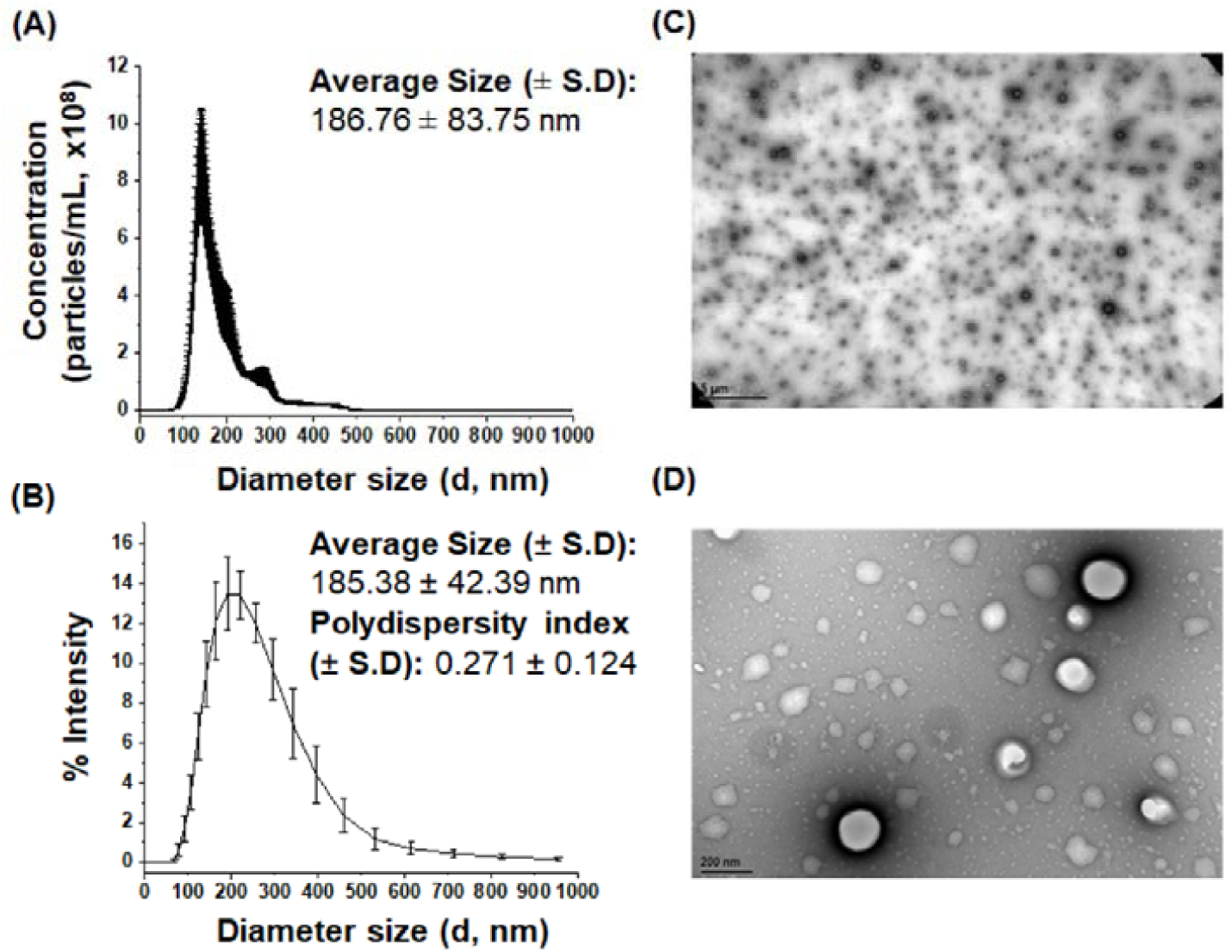
Characterization of CDDO-Me-NPs. The hydrodynamic diameter CDDO-Me-NPs and polydispersity index were measured via **(A)** nanosight nanotracking analysis and **(B)** dynamic light scattering. These graphs are representative of 3-8 separate preparations of NPs and the average size was determined ± 1 standard deviation. CDDO-Me-NPs were adsorbed onto copper 400 mesh TEM grids were imaged via conventional TEM at **(C)** 5000x magnification (scale bar = 5μm) and **(D)** 100,000x magnification (scale bar = 200 nm).

Replacement of the cholesteryl ester/triacylglycerol core of plasma LDL with OA, as described by the reconstitution method, yields LDL particles that are uniformly loaded with OA^21^. Herein we generated NPs that were fluorescent by co-loading with the lipophilic dye, DiD. Importantly, as per previous reported preparations, un-detectable amounts of cholesterol remain in the LDL nanoparticles reconstituted with OA^21^. Compositional analysis shows that, on average the fluorescently labelled LDL-DiD-OA NPs contained an estimated 57 ± 3 molecules of DiD fluorophore per particle. According to previous preparations each LDL is typically associated with 2000 molecules of unesterified OA^21^. The zeta potential for these particles was −36.7 ± 0.9 mV. Currently we were unable to perform DLS size measurements as the concentration of dye interferes with the light scattering process.

### 3.2 Activation of Nrf2 by CDDO-Me-NPs

Utilizing an *in vitro* Nrf2 activation assay^23, 24^, we confirmed that CDDO-Me-NPs have a delayed release of native CDDO-Me over 24 hours (Fig. 2A). In this assay, we show that CDDO-Me, a well-known Nrf2 activator, activates the transcription of the canonical down-stream target of Nrf2, NQO1 in a dose-dependent manner (10-400 nM). This is quantified as thresholded fluorescence intensity corresponding to GFP fluorescence per cell with an average of 200 cells/well and 6 wells measured per condition (representative images shown in Fig. 2B). In contrast, neither 1%DMSO nor polymer/DiD alone activate NQO1 transcription over 24 hours (Fig. 2). On the other hand, CDDO-Me-NPs activate NQO1 transcription via Nrf2 albeit to a lesser extent in the same time period. Indeed at the equimolar dose of 10nM CDDO-Me-NPs did not release sufficient CDDO-Me to induce a significantly different degree of NQO1 transcription. This indicates that the NPs exhibit delayed release of CDDO-Me over the 24 hours period.

**Figure 2.**
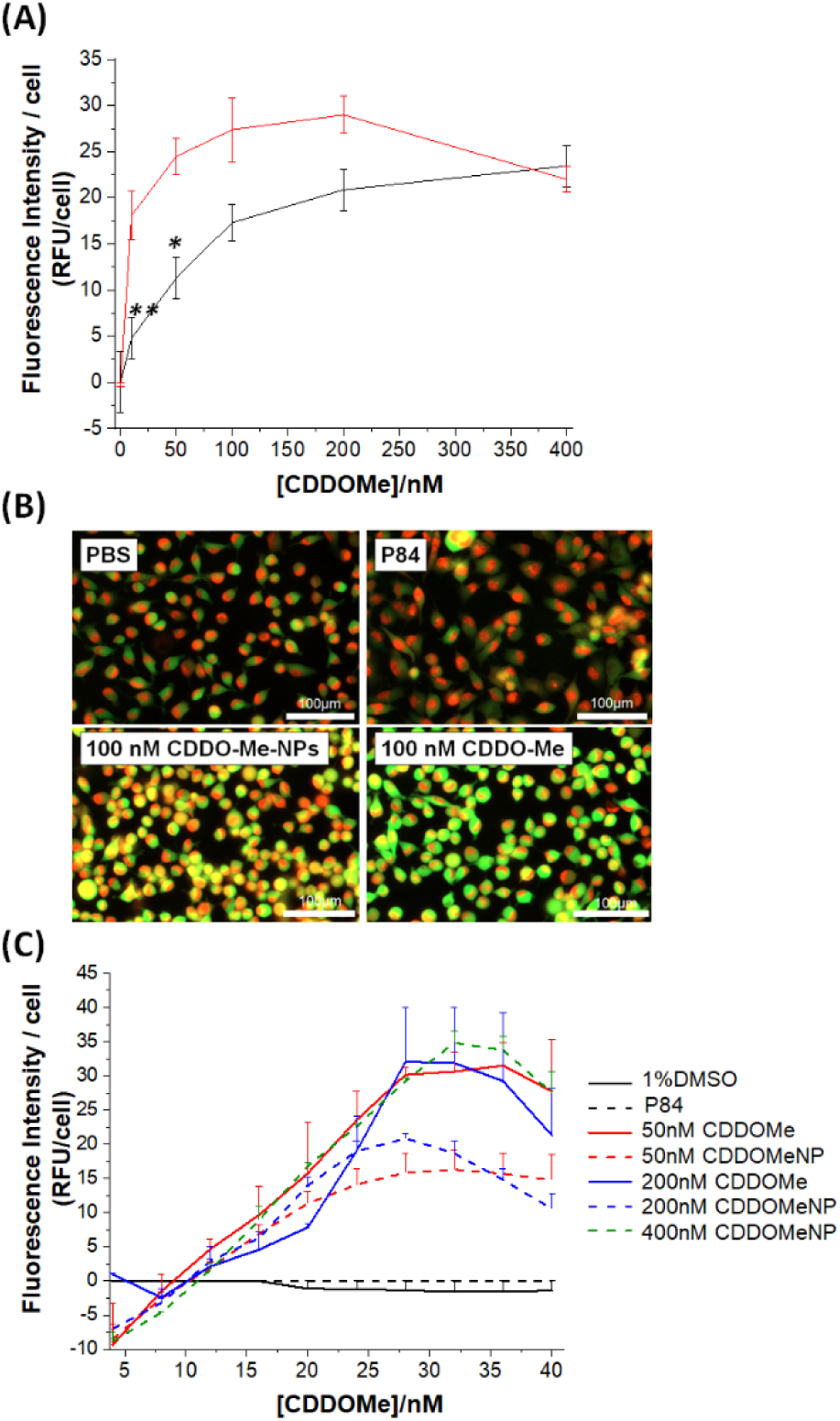
Activation of Nrf2 by CDDO-Me-NPs. **(A)** Dose-dependent activation of NQO1 transcription by CDDOMe and CDDOMeNPs. H1299 cells were treated with either 1% DMSO, P84 polymer (at equivalent concentrations as present in most concentrated NP preparations), CDDO-Me (10-400nm), or CDDO-Me-NPs (10-400nM) for 24 hours. GFP fluorescence intensity per cell was quantified with an average of 200 cells counted per measurement. * p<0.05, ** p<0.01, 2-way ANOVA with Tukey correction comparing CDDOMe vs. CDDOMeNP ± St.Dev, n=4-5 independent experimental data points. **(B)** Representative images of GFP fluorescence due to treatments to induce Nrf2 activation in H1299 cells after 24 hours as collected by the Cytation 5 plate reader. Scale bar is 100μm. **(C)** Time-dependent activation of NQO1 transcription by CDDOMe and CDDOMeNPs. H1299 cells were treated with either 1% DMSO, P84 polymer (at equivalent concentrations as present in most concentrated NP preparations), CDDO-Me (solid line, 50-200nm), or CDDO-Me-NPs (dashed line, 50-400nM) for up to 40 hours. GFP fluorescence intensity per cell was quantified with an average of 200 cells counted per measurement. ± SEM, n=2 independent experimental data points.

We then examined the activation of Nrf2 by CDDO-Me and CDDO-Me-NPs over the course of 40 hours at varying concentrations (Fig. 2C). Herein we found that NQO1 transcription is sustained between 24-40 hr. We additionally confirmed that higher concentrations of CDDO-Me-NPs are required (up to 400nM) to sustain NQO1 transcription at similar levels to un-encapsulated CDDO-Me.

### 3.3 Inhibition of VSMC migration by CDDO-Me-NPs

We next investigated whether CDDO-Me-NPs could inhibit VSMC migration. In our previous work we showed that Nrf2 activation inhibits migration^30^. Herein, using the scratch wound assay with murine aortic smooth muscle cells, we show that CDDO-Me can dose-dependently inhibit PDGF-BB-stimulated SMC migration over 48 hrs (Fig. 3). We additionally show that CDDO-Me-NPs significantly inhibit SMC migration over 48 hr, although higher concentrations are required to achieve a similar degree of inhibition. This is consistent with the delayed release of CDDO-Me.

**Figure 3.**
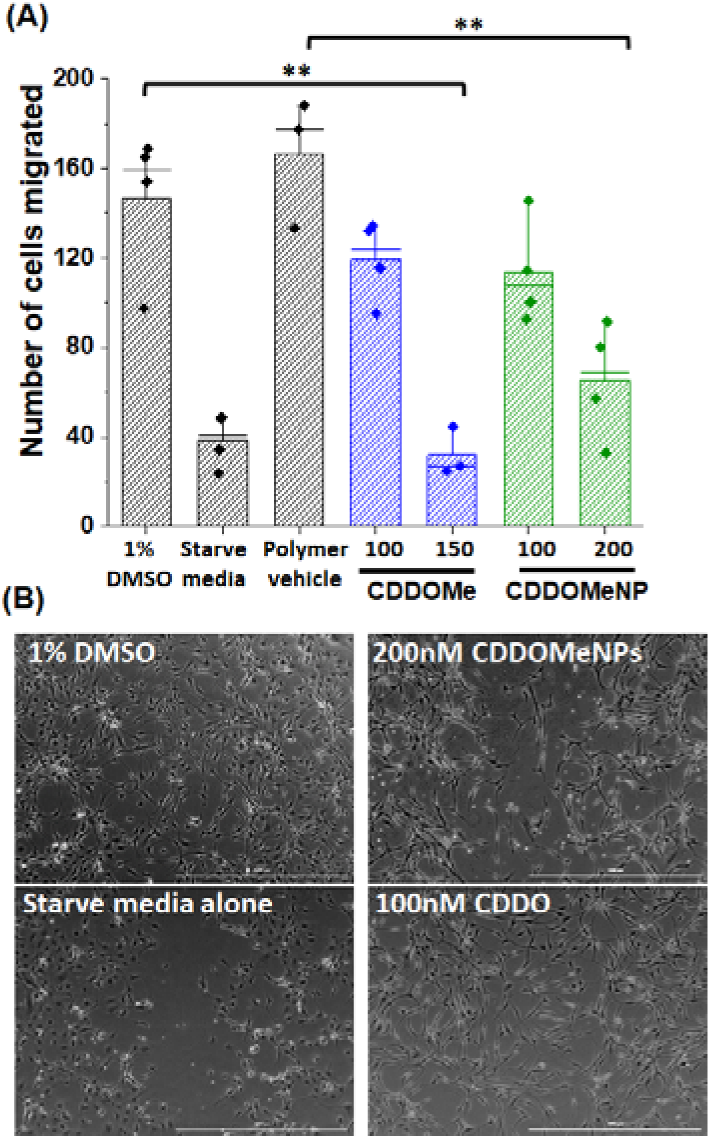
CDDO-Me-NPs and VSMC migration. **(A)** VSMCs were synchronized overnight and then either stimulated with PDGF-BB (25ng/mL) for 48 hours or kept in starvation media (no PDGF-BB). They were incubated in the presence or absence of treatments: 1% DMSO, P84 polymer (at equivalent concentrations as present in most concentrated NP preparations), CDDO-Me (100, 150nm), or CDDO-Me-NPs (100, 200nM). **(B)** Representative images of migration over 48 hours into the scratch wound. Phase contrast images of PDGF-treated (+) or PDGF-starved (-) cells were taken at 0 and 48 hr and the number of cells that migrated into the scratch wound were determined and represented graphically. Data represented as ± 1 standard deviation (n = 3-4 independent experiments in triplicate). Data was analysed via a two-way ANOVA with tukey correction, ** p<0.01.

### 3.4 Time-dependent internalization of CDDO-Me-NPs by RAW macrophages

Based on the critical role of macrophages in driving inflammation-mediated progression of atherosclerosis, we utilized a murine macrophage cell line, RAW 264.7 macrophages to examine the association and internalization of CDDO-Me-NPs with macrophages. CDDO-Me-NPs were labelled with DiD lipophilic dye and then incubated with RAW macrophages over 18 hours (Fig. 4A). As observed in Fig. 4A, thresholded fluorescence area/cell increased over 8 hours after which they plateaued. Fig 4B shows the fluorescence intensity/cell at the 18 hour time point after incubation with fluorescent CDDO-MeNPs, whilst Fig. 4C shows a representative image of RAW macrophages in the presence of CDDO-Me-NPs after 18 hours. To confirm that this signal was due to internalization we performed a temperature dependent assay whereby fluorescence CDDO-Me-NPs were incubated with RAW macrophages over 18 hr at either 4 or 37°C (Fig. 4D). At 4°C, energy-dependent internalization processes are inhibited, therefore the signal derived from RAW macrophages incubated at this temperature is CDDO-Me-NPs that are associated with the external cell surface. This assay confirms that a significant amount of the signal is due to internalization of CDDO-Me-NPs by RAW macrophages.

**Figure 4.**
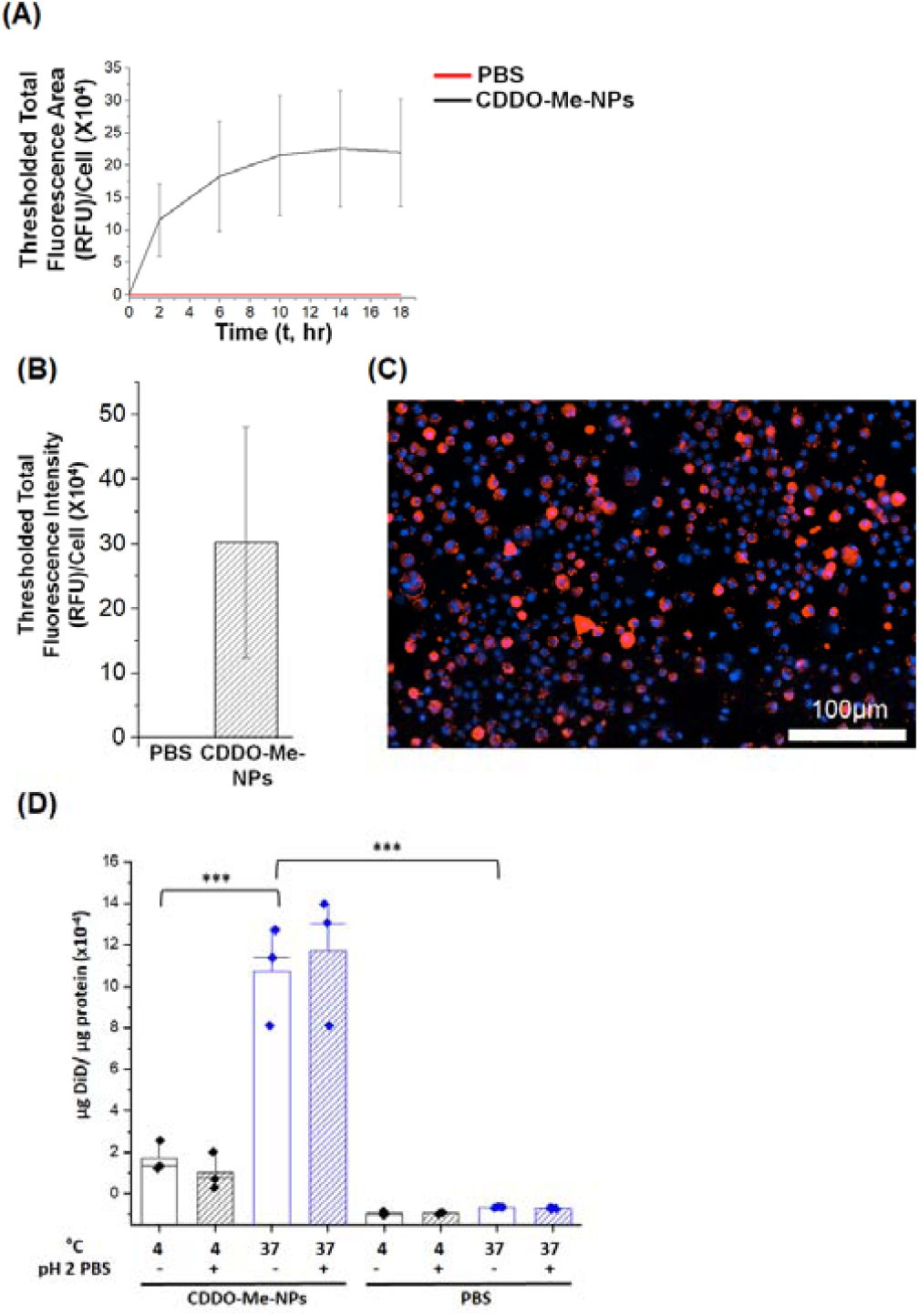
Association of CDDO-Me-NPs with RAW macrophages. RAW macrophages stained with Hoechst dye (10μg/mL, 30 minutes) were treated with fluorescent CDDO-Me-NPs (0.1mg/mL final, 20% PBS) over the course of 18 hrs. **(A)** Fluorescent images of RAW macrophages were taken at 4 hour intervals over 18 hours and thresholded fluorescence area was quantified per cell, an average of 800 cells were counted per well. Data represented as ±SEM (n=2 independent experiments in sextuplicate). **(B)** Fluorescent images of RAW macrophages were taken at 18 hours and thresholded fluorescence intensity was quantified per cell, an average of 800 cells were counted per well. Data represented as ±SEM (n=3 independent experiments in sextuplicate). Data is non-significant. **(C)** A representative image of macrophage-associated fluorescent CDDO-Me-NPs. Scale bar is 100μm. **(D)** Temperature-dependent internalization of CDDOMeNPs at 18 hr by RAW macrophages. Data represented as ±St.Dev (n=3 independent experiments in triplicate). ***, p<0.001.

### 3.5 Localization of CDDO-Me-NPs in atherosclerotic plaque in vivo

To assess whether our nanoparticles accumulated in atherosclerotic lesions we injected athero-prone mice with 2.5mg/kg of CDDO-Me-NPs. This was an equivalent dose of 25μg/kg of DiD fluorescent dye. 24 hrs following the injection we sacrificed these animals and excised the hearts for histology. We quantified thresholded fluorescence intensity normalized to lesion area (Fig. 5A) or lipid area (Fig. 5B, as determined by ORO staining) in the atherosclerotic plaque of the aortic sinus of both LDLr^-/-^ and ApoE^-/-^ mice. The increase in fluorescence intensity per lesion area and per lipid area was highly significant in both strains of athero-prone mice. Fig. 5C shows representative fluorescence images of the aortic sinus region of ApoE^-/-^ animals injected either with PBS or with 2.5mg/kg of CDDO-Me-NPs. This data confirms that CDDO-Me-NPs localize in atherosclerotic plaque.

**Figure 5.**
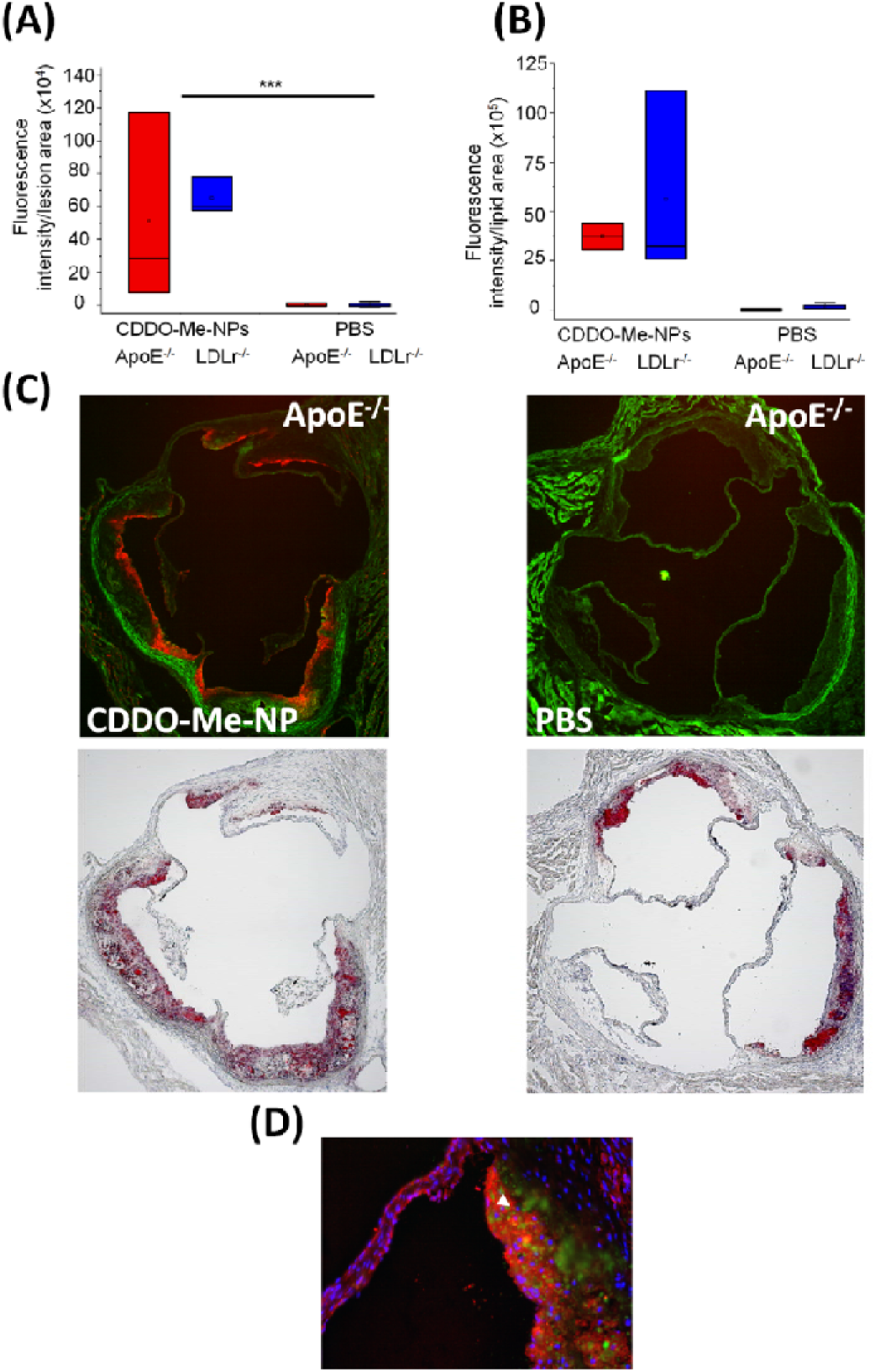
CDDO-Me-NPs localize in atherosclerotic plaque of the aortic sinus in athero-prone mice. 2.5mg/kg of fluorescent CDDO-Me-NPs in 10mM PBS or PBS were intravenously injected into 15 week high fat diet-fed ApoE^-/-^ or LDLr^-/-^ mice (18-19 weeks old). 24 hours following the injection, the heart was excised and prepared for histological cryo-sectioning. **(A)** Quantification of thresholded fluorescence intensity per lesion area. Data is represented as ±1 standard deviation, *** is p<0.001 between the CDDO-Me-NP injected and PBS injected animals (n = 4-5 animals, with 2-5 replicate tissue sections analyzed) **(B)** Quantification of thresholded fluorescence intensity per lipid area. Data is represented as ±1 standard deviation, *** is p<0.001 between the CDDO-Me-NP injected and PBS injected animals (n = 4-5 animals, with 2-5 replicate tissue sections analyzed). **(C)** Representative 5x magnification images of aortic sinus from ApoE^-/-^ animals injected with either LDL-OA-NPs or PBS. **(D)** Localization of NPs (green) with macrophages (CD11b-stained, red) in atherosclerotic plaque of LDLr^-/-^ mice.

### 3.6 Localization of LDL-OA-NPs in atherosclerotic plaque in vivo

In a similar manner, we assessed whether fluorescent LDL-OA-NPs accumulated in atherosclerotic lesions of athero-prone mice. These LDL-OA-NPs were co-loaded with DiD fluorescent dye. We injected the equivalent of 400μg/kg of DiD and then 24 hrs following the injection we sacrificed this animals and excised the hearts for histology. Once again, the increase in fluorescence intensity normalized to either lesion area or lipid area was highly significant in both strains of athero-prone mice (Fig. 6A, B). Fig. 6C shows representative fluorescence images of the aortic sinus region of ApoE^-/-^ animals injected either with PBS or with LDL-OA-NPs. Finally, we performed light sheet fluorescence microscopy of immune-stained, cleared and agarose-embedded aortic arches of LDLr^-/-^ animals that had either been injected with LDL-OA-NPs or PBS (Fig. 7A, B). In Fig. 7A(i) and 7B(i) clear localization of DiD fluorescence can be observed in atherosclerotic plaque in the aortic arch, which is absent in PBS-injected animals (Fig. 7A(ii), 7B(ii)). Colocalization with macrophages can additionally be observed in Fig. 7A where aorta were immuno-stained for CD68. This data confirms that LDL-OA-NPs localize in atherosclerotic plaque.

**Figure 6.**
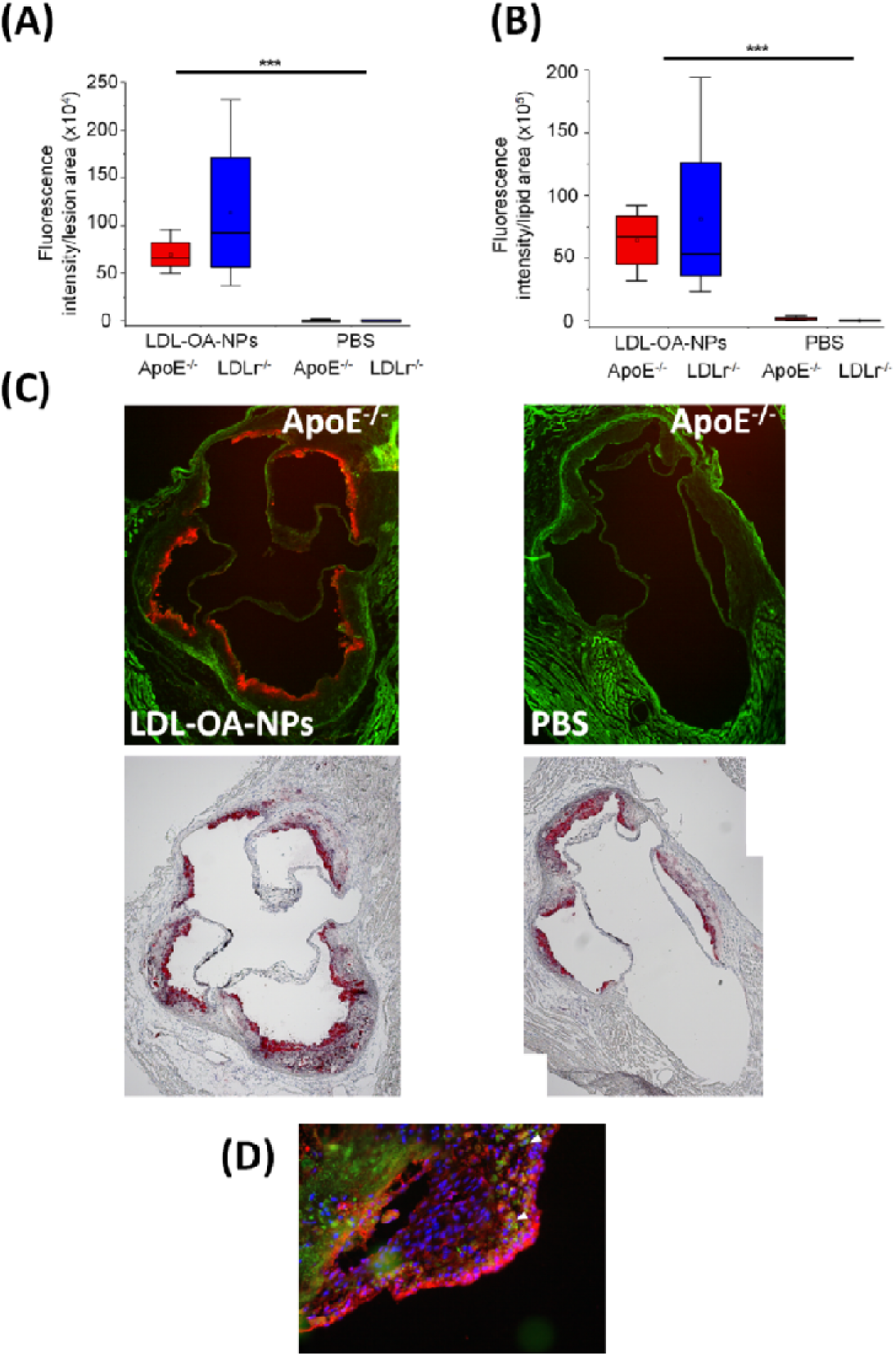
LDL-OA-NPs localize in atherosclerotic plaque of the aortic sinus in athero-prone mice. Fluorescent LDL-OA-NPs (400μg/kg of DiD) in 10mM PBS or PBS were intravenously injected into 19 week high fat diet-fed ApoE^-/-^ or LDLr^-/-^ mice (23-24 weeks old). 24 hours following the injection, the heart was excised and prepared for histological cryo-sectioning. **(A)** Quantification of thresholded fluorescence intensity per lesion area. Data is represented as ±1 standard deviation, *** is p<0.001 between the LDL-OA-NP injected and PBS injected animals (n = 4-5 animals, with 2-5 replicate tissue sections analyzed) **(B)** Quantification of thresholded fluorescence intensity per lipid area. Data is represented as ±1 standard deviation, *** is p<0.001 between the LDL-OA-NP injected and PBS injected animals (n = 4-5 animals, with 2-5 replicate tissue sections analyzed). **(C)** Representative 5x magnification images of aortic sinus from ApoE^-/-^ animals injected with either LDL-OA-NPs or PBS. **(D)** Localization of NPs (green) with macrophages (CD11b-stained, red) in atherosclerotic plaque of LDLr^-/-^ mice.

**Figure 7.**
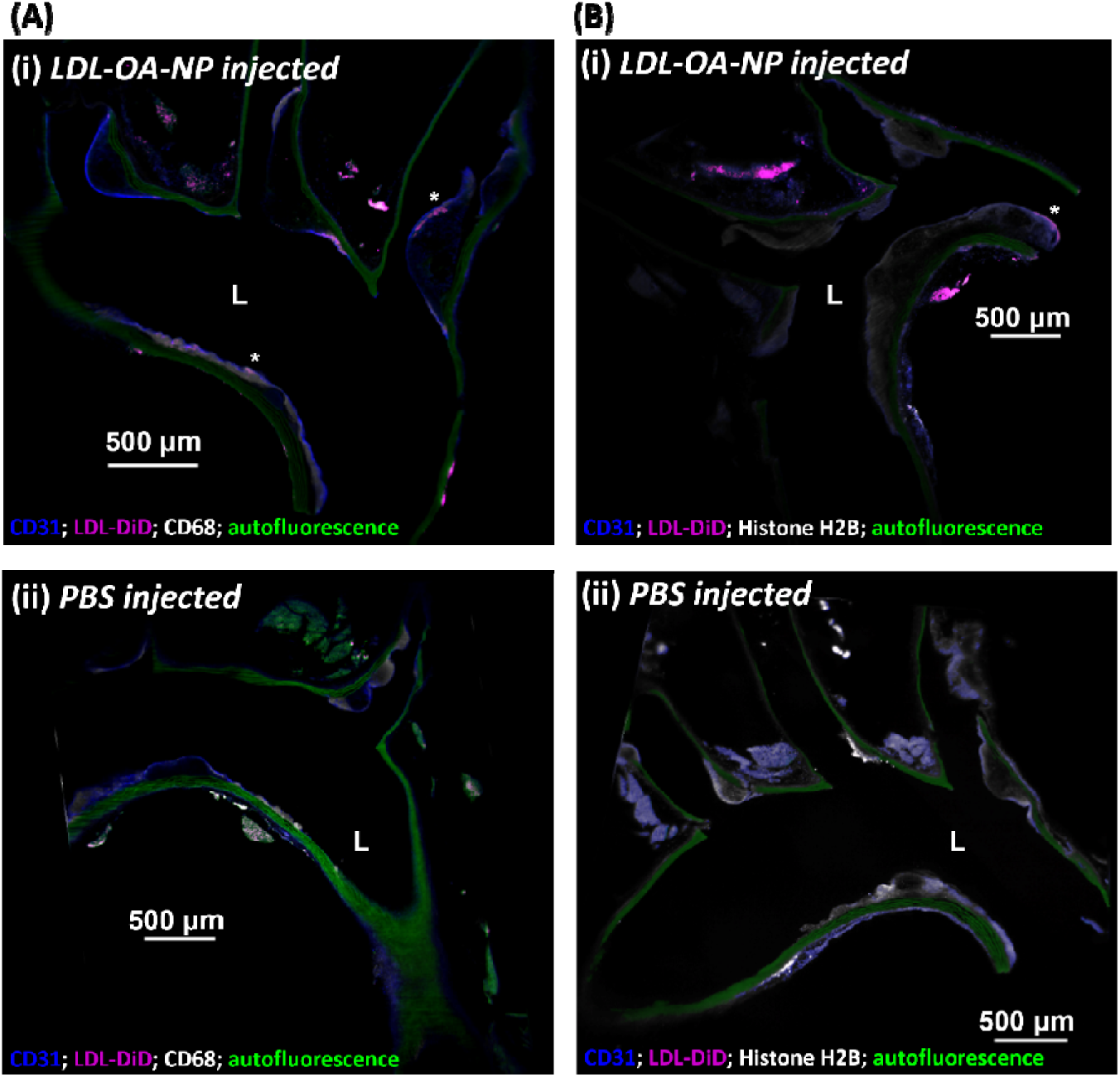
LDL-OA-NPs localize in atherosclerotic plaque in aortic arch of LDLr^-/-^ mice. Light sheet fluorescence microscopy rendering of the aortic arch. Representative slice views of the aortic arch of HFD fed LDLr^-/-^ animals. Aorta underwent processing for LSFM following methods according to previous publications^28, 29, 31^ and were immuno-stained for CD31 (blue) and either CD68 (white) or Histone H2B (white). Fluorescence due to LDL-DiD-OA-NPs is shown in magenta as well as auto-fluorescence due to excitation with the 488 nm laser (green). (A) LDL-DiD-OA-NPs co-localize with macrophages in atherosclerotic plaque. Asterisks indicate locations showing uptake of LDL-DiD-NPs in aorta. (B) LDL-DiD-OA-NPs localize in highly cellular regions of atherosclerotic plaque. Asterisks indicate locations showing uptake of LDL-DiD-NPs in aorta.

## 4. Discussion

The present study sought to use NPs to target atherosclerotic plaque through passive targeting means in 15-19 week old ApoE^-/-^ and LDLr^-/-^ mice fed with an HFD. Herein we generated two types of NPs: 1. Polymeric NPs encapsulating the NRf2 activator drug, CDDO-Methyl, and 2. LDL-like NPs encapsulating OA. We primarily found that both NPs accumulate significantly in atherosclerotic plaque of two widely used murine models of atherosclerosis.

### 4.1 CDDO-Me-NPs

The CDDO-Me-NPs were generated by the FNP technique. This technique is highly useful due to the rapid formulation as well as the high cargo loading^19^. At the time of writing there appears to be only one other literature report of a polymeric formulation of CDDO-Me with PLGA, where the drug loading capacity 2.9 ± 0.2%, after being generated by the solvent displacement method^32^. With our method our NPs achieve a drug loading capacity of (35±7)%. This speaks to the value of FNP in terms of the capacity of drug that can be included per mg of nanoparticles. The CDDO-Me-NPs were designed to be close to 200nm as a result of the literature showing that for uptake of NPs by macrophages this size is optimal^33, 34^. Previous literature has also shown that size is an important determinant for accumulation in atherosclerotic lesions, with a 200nm sized liposome providing the best accumulation^33^. This has also been shown by Ogawa and colleagues^35^. In this current study we observed a time-dependent association of polymeric CDDO-Me-NPs with murine RAW macrophages, which plateaued after 8 hours. Moreover, *in vivo* we found evidence via immunofluorescence of macrophages with CD11b staining of association of both CDDO-Me-NPs and LDL-OA-NPs with atherosclerotic plaque macrophages.

Their accumulation is consistent with the literature which shows that NPs can also accumulate selectively in atherosclerotic plaque simply due to the enhanced permeation and retention (EPR) effect^36, 37^. NPs take advantage of a dysfunctional endothelium with larger inter-endothelial junctions to accumulate in sites of vascular inflammation^36, 37^. Early atherosclerotic events have been shown to be associated with more severe endothelial barrier disruption, whilst advanced stage disease progressions are associated with improved endothelial barrier permeability^36^. Previous pre-clinical studies have successfully integrated NPs into sites of atherosclerotic plaque, some of which were able to inhibit plaque formation and decreased the number of atherosclerotic lesions in mouse models of atherosclerosis^33, 37-41^. Macrophage-mediated uptake of NPs *in vivo* is also likely to reflect phagocytosis due to the presence of the NP-protein corona that plays a role in NP-cell interactions^42^.

It was beyond the scope of this study to assess the therapeutic effects of intra-plaque delivered Nrf2 activator drugs such as CDDO-Me, which will instead be assessed in future work. The present study did show that our NP formulation exhibited a delayed release of CDDO-Me to active Nrf2 *in vitro*. Moreover, in the literature it has previously been shown that activation of Nrf2 in LDLr^-/-^ with tBHQ can have atheroprotective effects based on the alleviation of oxidative stress and limitation of inflammation^10^. It is known that Nrf2 activation inhibits VSMC migration^30^. Consistent with past work that has reduced inflammation at sites of vascular injury, CDDO-Me inhibited PDGF-BB-stimulated SMC migration over 24 hrs in a dose-dependent manner; the non-significant but dose-dependent trend of CDDO-Me-NPs is also consistent with the delayed drug release, as stated previously. Despite future work being needed to investigate the inhibition of migration at longer time points, our data indicate that the sites of atherosclerotic plaque were successfully targeted by both CDDO-Me-NPs and LDL-OA-NPs. Monocyte-turned-macrophage aggregation in the endothelium contributes to the non-resolving inflammatory response of atherosclerosis. Collectively, the successful localization of the NPs and macrophage uptake of the CDDO-Me-NPs in addition to the release of CDDO-Me and inhibition of VSMC propose a promising therapeutic strategy to reduce atherosclerosis, by targeting specific sites of injury.

### 4.2 LDL-OA-NPs

In recent years, the dietary benefits of omega-3 polyunsaturated fatty acids (ω-3 PUFAs) and oleic acid have been heralded for improving cardiovascular health^22^. Therapeutic intravenous administration of oleic acid or ω-3 PUFAs is not a realistic option due to poor aqueous solubility and propensity to form lipid emboli^43, 44^. The concept of utilizing LDL as a drug-delivery vehicle was first proposed over three decades ago^45-48^. Utilizing lipoproteins as a delivery vehicle for oleic acid takes advantage of the physiological function of lipoprotein particles to transport lipids in plasma^49^. Moreover, LDL particles represent an attractive delivery vehicle in particular for atherosclerosis, where the uptake of these particles into plaque represents an intrinsic feature of atherogenesis^50^. In the present study, oleic acid was uniformly incorporated into low-density lipoprotein (LDL) nanoparticles (LDL-OA-NPs) as previously reported^21^ with the addition of the lipophilic dye, DiD, in order to visualize their distribution in athero-prone mice. Being a natural lipid, OD readily incorporates into LDL through the reconstitution process. The physicochemical and stability properties of this novel nanoparticle were evaluated; in addition, the selectivity of the LDL-OA-NPs to accumulate in atherosclerotic plaque was evaluated.

The reconstitution method of generation LDL-OA-NPs is also highly efficient and represents a facile method to establish high cargo loading of fatty acids and ω-3 PUFAs into LDL particles. According to previous preparations each LDL is typically associated with 2000 molecules of unesterified OA^21^, which is an excellent comparison to the number of cholesteryl ester molecules typically associated with native LDL particles (∼1500). In previous work in the Corbin lab we have shown that the reconstitution method of generating LDL-like NPs results in NPs that retain nearly all of the physicochemical properties and biological activities of their native counterparts^20, 21^. Particle size, morphology and apoB-100 protein conformation were all comparable with that seen in native LDL. This is yet to be definitely confirmed for the fluorescent counterpart of LDL-OA-NPs as presented in this work.

LDL uptake and retention in the intimal layer of the vessel wall is an intrinsic aspect of atherogenesis^50^. The initial fatty streak phase of atherosclerosis begins with the retention of LDL particles resulting in a dysfunctional endothelium. As atherosclerotic plaque progress, modified LDL is recognised by infiltrating monocytes leading to their cellular internalization and the formation of macrophage foam cells. It is this natural retention and internalization mechanisms that led to the interest in using reconstituted LDL-NPs as delivery vehicles for anti-inflammatory, antioxidant fatty acids and ω-3 PUFAs. Indeed, in the present study we unequivocally demonstrate a selective accumulation of LDL-OA-NPs in atherosclerotic plaque in two animal models of atherosclerosis. We additionally show that these LDL-OA-NPs associated with macrophages in atherosclerotic plaque. In future work it will be important to assess whether a therapeutic benefit is conferred by a treatment regimen of administration of LDL-OA-NPs or LDL-DHA-NPs in inhibiting atherosclerotic plaque progression. Furthermore, it remains to be seen whether intravenous administration of reconstituted LDL-NPs will worsen atherosclerotic burden by providing greater amounts of oxidized LDL, resulting in for instance endothelial cell activation. The susceptibility of reconstituted LDL-NPs to oxidative modification and whether this occurs *in vivo* must be determined. On the other hand, the presence of fatty acids, such as oleic acid, or ω-3 PUFAs may act to render the reconstituted LDL-NPs less susceptible to oxidation and act to reduce oxidation and inflammation within the vascular wall.

## 5. Conclusion

Herein we generated two types of NPs: 1. Polymeric NPs encapsulating the Nrf2 activator drug, CDDO-Methyl and 2. LDL-like NPs encapsulating the anti-inflammatory compound oleic acid. We show that CDDO-Me-NPs exhibit a delayed release of native CDDO-Me to activate Nrf2 *in vitro*. Furthermore we show a dose-dependent effect upon VSMC migration with CDDO-Me-NPs. We then showed that CDDO-Me-NPs associate with RAW macrophages. Finally we showed that both CDDO-Me-NPs and LDL-OA-NPs accumulated selectively in atherosclerotic plaque in two murine models of atherosclerosis.

## Acknowledgements

We gratefully acknowledge Vicky Madden and Kristen White for assistance with conventional TEM. The Microscopy Services Laboratory, Department of Pathology and Laboratory Medicine, is supported in part by P30 CA016086 Cancer Center Core Support Grant to the UNC Lineberger Comprehensive Cancer Center. We also gratefully acknowledge Olesia Gololobova and Jacob Ramsey of the Nanomedicines Characterization Core Facility (NCore) at the Center for Nanotechnology in Drug Delivery (CNDD) at UNC School of Pharmacy for their assistance with Nanosight and HPLC UV-VIS analysis of nanoparticle samples.

